# Reduced variation in *Wolbachia* density of larval stages in comparison with adults of *Onchocerca volvulus:* implications for clinical outcome of infection?

**DOI:** 10.1101/458034

**Authors:** Samuel Armoo, Stephen R Doyle, Shannon M Hedtke, Gilles A Adjami, Daniel A Boakye, Annette C Kuesel, Mike Y Osei-Atweneboana, Warwick N Grant

**Affiliations:** Biomedical and Public Health Research Unit, Council for Scientific and Industrial Research - Water Research Institute, GA038, Council Close, Accra, Ghana; Wellcome Sanger Institute, Hinxton, Cambridgeshire, CB10 1SA, United Kingdom; Department of Animal, Plant and Soil Sciences, School of Life Sciences, La Trobe University, Bundoora 3083, Victoria, Australia; The Expanded Special Project for Elimination of Neglected Tropical Diseases, Ouagadougou, Burkina Faso; UNICEF/ UNDP/ World Bank/ WHO Special Programme for Research and Training in Tropical Disease, WHO, Geneva 1211, Switzerland

## Abstract

**Background:** *Wolbachia* are important endosymbionts of filarial parasites. The *Wolbachia* of *Onchocerca volvulus*, the filarial pathogen responsible for the human disease onchocerciasis, is implicated in the immunopathology of the disease and may be associated with disease severity dependent on the density of *Wolbachia*. However, little is known in regards to the density and heterogeneity of *Wolbachia* in microfilariae, the life stage that is thought to be responsible for the pathology.

**Results:** We used a real-time qPCR relative copy number assay to estimate the number of *Wolbachia* genome(s) per nuclear genome of skin microfilariae (Mf), vector L1 and iL3, and nodulectomy adult male and female *O. volvulus* worms sampled in Ghana and the Democratic Republic of the Congo. Relatively low median *Wolbachia* copy numbers and variation was observed in the Mf and vector stages, in contrast to significantly higher median and more variable *Wolbachia* copy number from the iL3 stage to the adult worm stages.

**Conclusions:** This study provides the first insight into variation in *Wolbachia* density between the major life stages of the parasite. The relatively invariant ratios observed for Mf and vector stages is in strong contrast to the high degree of variability of *Wolbachia* to nuclear ratios in adults and may indicate that the mutualistic relationship between the nematode and *Wolbachia* in these earlier stages is regulated differently, and certainly more stringently, than the relationship in adults.

## Background

The human disease onchocerciasis, caused by the filarial nematode *Onchocerca volvulus*, remains a significant public health problem in Sub-Saharan Africa. The immunopathology associated with the disease, which includes dermatitis and keratitis, has been linked to a *Wolbachia* endosymbiont that is immunologically recognised by the host upon the death of microfilaria [1, 2, 3]. Evidence of this includes the initiation of keratitis by *Wolbachia* antigens in a murine model [1], and a significantly higher proportion of *Wolbachia* DNA in the sera from patients following treatment with either diethyl carbamazine or ivermectin [3]. The association between *Wolbachia* and pathology has been extended to suggest that *Wolbachia* density in adult worms is positively correlated with the incidence of blindness [4]. This hypothesis was based on measurements that compared *Wolbachia* copy number from “forest ecotype” and “savannah ecotype” *O. volvulus* that are associated with low blindness rates and low *Wolbachia* density, or higher blindness and higher *Wolbachia* densities, respectively. However, using larger and geographically diverse cohort of adult *O. volvulus* samples, we recently demonstrated using qPCR that the *Wolbachia* copy number ranges over one thousand fold in adult worms, and differs within and between sampling locations independent of the “forest” and “savannah” ecotype [5]. These results, therefore, did not support the original hypothesis that *Wolbachia* copy number in adult worms is associated with ocular pathology.

Much of the evidence supporting a leading role for *Wolbachia* as a driver of onchocerciasis pathology comes from *in vitro* experiments using adult worm extracts. However, both the chronic inflammation of the skin and eyes and the acute Mazzotti reaction pathology are due to release of material from dying microfilaria. Any correlation between *Wolbachia* and pathology (including a correlation with *Wolbachia* density) must, therefore, be supported by appropriate data on microfilarial *Wolbachia*. We report here a comparison of *Wolbachia* density, measured by qPCR, in skin microfilaria, infective larvae and adult parasites, and show that the density in skin microfilaria is low and relatively constant, in contrast to a higher and more variable *Wolbachia* density in adult worms. This observation further supports a view that variation in pathology is unlikely to be influenced strongly by variation in *Wolbachia* density in adult worms.

## Results

The life-stage-specific distribution of *Wolbachia* copy numbers in field isolates of *O. volvulus* is presented in Figure 1. The Mf and larval stages were characterised by relatively low *Wolbachia* copy numbers ratios and low variation between samples. The median *Wolbachia* copy number ratio for Mf was 0.18 (range = 0.07 to 0.54; Table 1), whereas the median for the L1 stage was 0.15 (range = 0.04 to 0.53; Table 1). With regards to the infective larval stage, the median *Wolbachia* copy number ratio was 0.35 (range = 0.04 to 0.89; Table 1). Pairwise Wilcoxon rank sum tests between life cycle stages (Table 2) showed no significant differences in the distribution of *Wolbachia* copy number ratios between the Mf and the iL3 stages (P = 0.1705); Mf and L1 stages (*P* = 0.7047); and the L1 and iL3 stages (*P* = 0.3286).

There was a significant increase in the median *Wolbachia* copy number from the iL3 stage to the adult stage for both males (approximately 18-fold increase; Figure 1; Table 2; *P* = 0.0035), and females (about 15-fold increase; Figure 1; Table 2; *P* = 0.0015), although the distributions of copy number overlapped across all life stages (i.e. Table 1: the upper limits of the copy number range for the larval stages overlapping with the lower limits of the copy number range for the adult stages). The median *Wolbachia* copy number ratios were similar between adult male and female *O. volvulus* isolates (Figure 1; Table 2; *P* = 0.8315), and their distributions did not differ stochastically based on a Wilcoxon rank sum test (or Mann-Witney U test; Table 2). However, there was significantly higher variation in *Wolbachia* copy number within adult male and female field isolates of *O. volvulus* compared to earlier life stages (Figure 1; Table 1; Table 2). *Wolbachia* copy numbers in adult male field isolates of *O. volvulus* ranged from 0.03 to 68.35, whereas those in adult female isolates ranged from 0.29 to 76.77 (Table 1).

**Figure 1:**
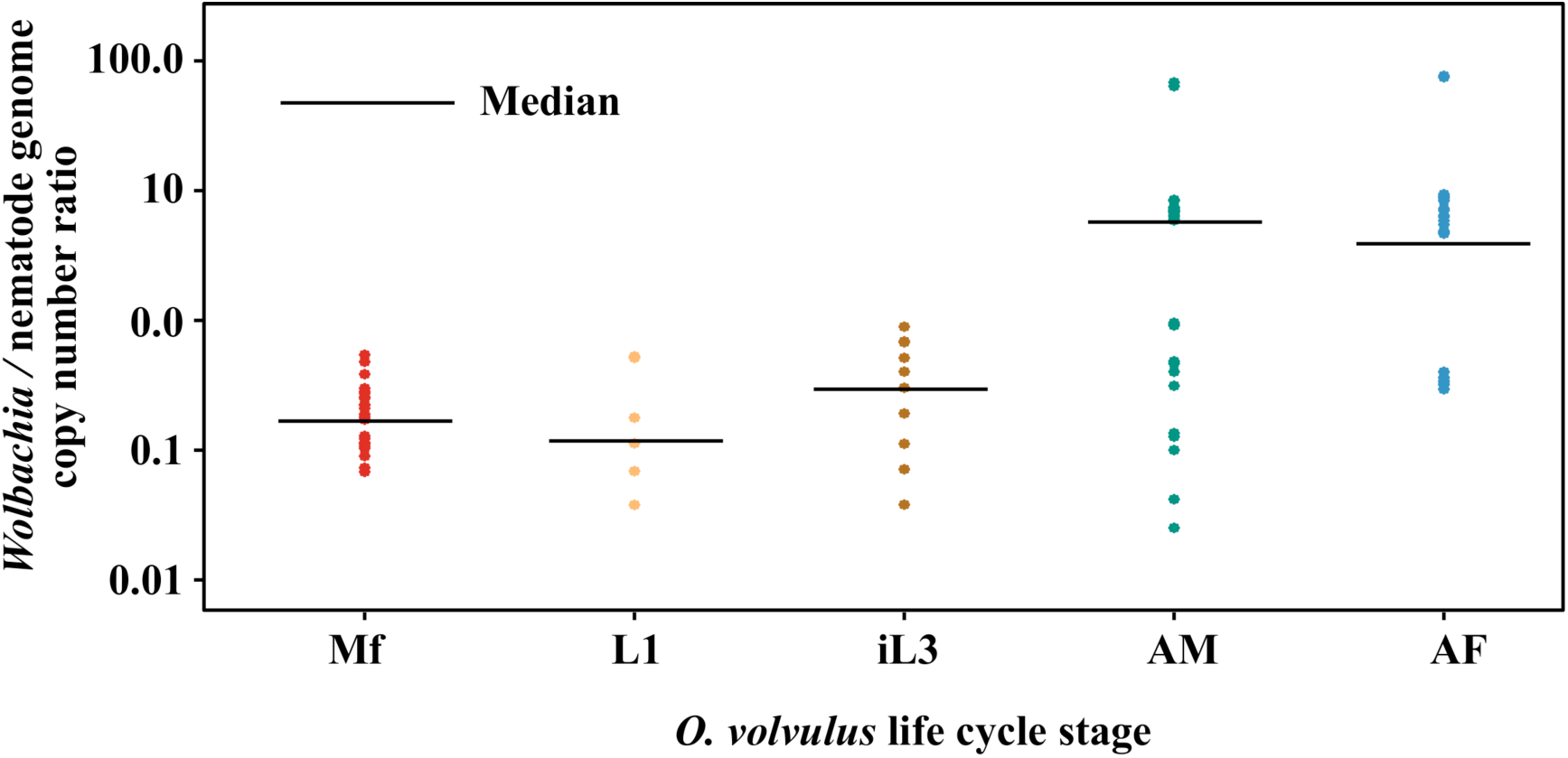
The density of *Wolbachia* (presented as *Wolbachia* / nematode genome copy number ratio) compared among different life stages of *Onchocerca volvulus.* The scatter plots show the distribution of *Wolbachia* copy numbers with each life stage. The black lines indicate the median values for each life stage. The plots are colour-coded to correspond with different life stages. Mf = microfilariae, L1 = first stage larvae, iL3 = infective stage three larvae, AM = adult males, AF = adult females.

**Table 1:** The median and range of *Wolbachia* copy number ratios of five *O. volvulus* life stages. Mf = microfilariae, L1 = first stage larvae, iL3 = infective stage three larvae, AM = adult males, AF = adult females

| *O. volvulus* life cycle stage | *Wolbachia* / nematode genome copy number ratio |  |
| --- | --- | --- |
|  | Median | Range |
| Mf | 0.18 | 0.07 to 0.54 |
| L1 | 0.15 | 0.04 to 0.53 |
| iL3 | 0.35 | 0.04 to 0.89 |
| AM | 6.40 | 0.03 to 68.35 |
| AF | 5.18 | 0.29 to 76.77 |

**Table 2:** P-values based on Wilcoxon rank sum tests for differences in *Wolbachia* density distributions between life stages. Mf = microfilariae, L1 = first stage larvae, iL3 = infective stage three larvae, AM = adult males, AF = adult females.

|  |  | *Onchocerca volvulus* life cycle stage |  |  |  |
| --- | --- | --- | --- | --- | --- |
|  |  | Mf | L1 | iL3 | AM |
| *Onchocerca volvulus*<br>life cycle stage | L1 | 0.7047 | - | - | - |
|  | iL3 | 0.1705 | 0.3286 | - | - |
|  | AM | 2.9e<sup>-6</sup>* | 0.0075* | 0.0035* | - |
|  | AF | 1.0e<sup>-11</sup>* | 0.0016* | 0.0015* | 0.8315 |

\* P < 0.05

## Discussion

In this study, we compared the relative copy number and heterogeneity of *Wolbachia* between larval and adult stages of *O. volvulus* from several sampling locations in central Ghana and the north-eastern region of the DRC. The homogeneous distribution of *Wolbachia* copy number in the Mf and larval stages of *O. volvulus* observed here is consistent with the findings recorded in *B*. *malayi* [6, 7, 8]. Electron microscopy studies have revealed that just a few cells of the Mf and larval stages of *B. malayi* contained sparsely distributed *Wolbachia* [8, 9].

In contrast to the homogeneous distribution of *Wolbachia* in Mf and larval stages, there was significantly greater variability in the copy number variation in both adult male and female worms, consistent with the findings of Armoo and colleagues [5], despite using a different set of adult male and female *O. volvulus* field isolates. Armoo and colleagues [5] used a qPCR assay to measure *Wolbachia* copy numbers in individual adult male and female worms sampled from four countries in West Africa (Togo, Ghana, Côte d’Ivoire and Mali), and detected significant within-population variation of *Wolbachia* copy numbers.

There are several possible explanations for variation in *Wolbachia* density between the life stages. Assuming that *Wolbachia* provide essential metabolites to the worm [10], the low and relatively homogeneous density of *Wolbachia* within all the larval stages could be explained by relatively low metabolic demands of early stage larvae, with a much smaller biomass and no reproductive tissues compared to an adult worm. Therefore the larval stages would likely require less from the mutualistic relationship than a large, reproductively active adult.

It is less clear how to account for the 100 – 1000 fold range in apparent *Wolbachia* density between individual adults, and the absence of this variability in microfilaria and larval stages. If the primary driver of the mutualistic relationship between worms and bacteria is metabolic [10, 11, 12], then this range implies either that adult worms can tolerate very variable levels of the metabolites they obtain from the bacteria without deleterious perturbation of their metabolism, or, that the metabolic rates of adult worms is variable. In the latter case (variation in adult worm metabolic rate), and *Wolbachia* density is coupled strongly to the adults’ metabolic rate, then variation in *Wolbachia* density implies that the bacteria are able to respond dynamically to the worms’ demand for metabolites. Preliminary genome wide association analysis of the *Wolbachia* and the worm nuclear genome did not reveal any obvious association of particular worm or bacterial genomes with high or low density, suggesting that the ratio of *Wolbachia* to nuclear genomes is not a genetically determined trait (Hedtke & Doyle, unpublished).

Although this study does not include data on the disease status of infected individuals from whom the Mf samples were obtained, the Mf analysed were from the north-east of the DRC, a region that is largely savannah and in which there is onchocerciasis associated blindness. The L1 and infective larvae samples were from central Ghana, which is also largely savannah and where there is also onchocerciasis-associated blindness. The low and homogeneous distribution of *Wolbachia* in the Mf and other larval stages may suggest that disease pathology or ecotype of the parasite may not be positively correlated with *Wolbachia* density in individual Mf.

## Conclusions

The similarity of the data presented in this report with data on *Wolbachia* densities in the related filarial pathogen, *B. malayi*, suggests that the regulation of *Wolbachia* copy number across life stages is evolutionary conserved and likely represents similar mutualistic strategies by the two parasites at similar stages of the life cycle. However, the mechanism(s) by which this regulation occurs, and the tolerance for significant variation, especially in the adult stages of the parasite, warrants further investigations. Understanding this regulation may be important in the context of anti-*Wolbachia* therapies that aim to kill adult parasites by depleting their *Wolbachia.* For example, an adult with very low *Wolbachia* density and hence presumably low demand for *Wolbachia* metabolites, may be less sensitive to depletion of the *Wolbachia.* Likewise, *Wolbachia* at low density may be in a less metabolically active state that may decrease their sensitivity to antibiotics that target metabolic processes such as protein translation (the target of doxycycline antibiotics). Furthermore, understanding the regulation of what may be an active, dynamic regulation of *Wolbachia* density may offer new insight into novel drug targets to disrupt the mutualistic relationship between the bacteria and its worm.

## Methods

### *O. volvulus* Samples

We used archival field parasite samples that were collected from central Ghana and north-eastern region of the Democratic Republic of the Congo. These field samples included 30 each of adult female and male *O. volvulus* worms, six first stage (L1) and ten infective third stage (iL3) larvae from three endemic communities in Ghana (Kyingakrom: latitude 8.0988; longitude −2.1090; Jagbengbendo: latitude 8.3342; longitude −0.1256; and Asubende: latitude 8.0171; longitude −0.9596). These samples from Ghana were collected from communities that were mainly found in the largely Savannah, and forest-Savannah transition zones. Therefore these parasites could be classified as “savannah ecotype”. In addition, we used 24 pools (each containing five individuals) of *O. volvulus* microfilariae (Mf) that were sampled from Lufu in the Democratic Republic of the Congo (latitude −5.68446; longitude 13.91585). Lufu is located within the Savannah, therefore could be classified as Savannah ecotype.

The archived L1 and iL3 larvae were previously isolated from the midgut and head region of blackfly vectors, respectively. The Biomedical and Public Health Research Unit of the Council for Scientific and Industrial Research (Accra, Ghana) carried out the field blackfly sampling. The same team isolated both adult male and female stages of *O. volvulus* after surgical removal of nodules from infected individuals in three endemic communities in central Ghana. The Mf were isolated from the skin snips (average weight of 1 mg) that had been taken from the iliac crest of infected individuals using the Holth-type corneoscleral punch. The Expanded Special Project for Elimination of Neglected Tropical Diseases, Ouagadougou, Burkina Faso, did sampling and archiving of Mf as a part of routine surveillance activities.

DNA was isolated from individual adult worms using the Bioline Isolate II genomic DNA extraction kit (Bioline, Sydney, Australia) following the manufacturer’s protocol. Individual L1 and iL3 worms and pools of MF were lysed in a 20 μl solution from a master mix of 98.5 μl of DirectPCR™ lysis reagent (MouseTail; Viagen Biotech, Los Angeles, USA) and 1.5 μl of 20 mg/ml Proteinase K stock (Roche Diagnostics GmbH, Mannheim, Germany) as previously described [13].

### Real-time qPCR *Wolbachia* Copy Number Assay

The relative *Wolbachia* copy number for each worm DNA extract was determined using a real-time qPCR assay designed and used previously [5]. In summary, the assay was designed based on two single copy genes in the *Wolbachia* and nematode genomes: the *Wolbachia* surface *(wsp)* gene (GenBank: HG810405.1) and the glutathione reductase *(gr*) gene (GenBank: Y11830.1) of the nematode. The sequence of the *wsp* targeting primers were forward: AACCGGGACAAAAAGAAGAG; reverse: CAGCAACCTACCAAAGATGGA, and that for the *gr* targeting primers were forward: GTGCGACGAAGAAGGATTTC; reverse: GCTTATGCTGTTTCGGGTTT.

Each qPCR reaction mixture (a total volume of 10 μl) consisted of 0.2 pmoles/μl of each primer, 2 μl of DNA and 5 μl of SsoAdvanced™ Universal SYBR^®^ Green Supermix (Bio-Rad Laboratories Inc., California, USA). All runs were performed in duplicate on the CFX 96 Real-Time PCR Detection System (Bio-Rad Laboratories Inc., California, USA), using the following thermal protocol: 95 °C for 2 min, followed by 40 cycles of 95°C for 5 sec, 53.8°C for 15 sec and 72°C for 15 sec. As a quality control measure, melt curves were generated at the end of each qPCR run to ensure specificity of primers.

The quantification cycles (Cq) of all qPCR runs were automatically generated by the CFX Manager Software v 3.1 (Bio-Rad Laboratories Inc., California, USA), and used to determine relative *Wolbachia* copy number of each sample as done elsewhere [5].

### Statistical Analyses

Data entry and re-formatting were performed using Microsoft Excel (2011). Wilcoxon rank sum tests (W) were performed using the R programming language, version 3.2.2 [14].

## Declarations

### Ethics approval and consent to participate

The DNA samples were obtained from archived parasite materials that were obtained from field studies by the Biomedical and Public Health Research Unit of the CSIR-Water Research Institute; and The Expanded Special Project for Elimination of Neglected Tropical Diseases. The Institutional Review Board of CSIR granted ethical clearance for CSIR team, and the WHO African Programme for Onchocerciasis Control covered the Expanded Special Project for Elimination of Neglected Tropical Diseases

### Consent for publication

All authors read, approved the final version of the manuscript and have consented to its submission to Parasites & Vectors for publication.

### Availability of data and material

All relevant data are included in the manuscript. Experimental materials are available from the corresponding author on request.

### Authors’ contributions

**Conceptualization**: Warwick N Grant, Stephen R Doyle, Samuel Armoo

**Investigation**: Samuel Armoo, Shannon M. Hedtke

**Methodology**: Samuel Armoo, Stephen R Doyle, Warwick N Grant

**Resources**: Mike Y Osei-Atweneboana, Daniel A Boakye, Gilles A Adjami, Stephen R Doyle, Warwick N Grant

**Data analysis**: Samuel Armoo

**Supervision**: Warwick N Grant, Stephen R Doyle, Annette C Kuesel, Mike Y Osei-Atwenebonana

**Writing** – original draft: Samuel Armoo.

**Writing** – review & editing: Samuel Armoo, Stephen R Doyle, Shannon M Hedtke, Gilles A Adjami, Daniel A Boakye, Annette C Kuesel, Mike Y Osei-Atweneboana, Warwick N Grant

## Acknowledgements

The authors would like to thank the field team of the Biomedical and Public Health Research Unit of the CSIR – Water Research Institute, and the Expanded Special Project for Elimination of Neglected Tropical Diseases.

